# Distinct geometrical landscapes distinguish between modes of tristability in gene regulatory networks

**DOI:** 10.1101/2025.10.11.681792

**Authors:** Atchuta Srinivas Duddu, Mohit Kumar Jolly, Archishman Raju

**Affiliations:** Department of Bioengineering, Indian Institute of Science, Bangalore, India; National Centre for Biological Sciences, Tata Institute of Fundamental Research, Bangalore, India

**Keywords:** cell fates, geometrical landscapes, gene regulatory motifs, EMT

## Abstract

Geometrical models have been recently used to construct landscapes for cell-fate decisions, inferred directly from experimental data. However, such quantitative cell-fate data is available for a few systems only; instead, gene regulatory networks dynamics have been studied for a broader set of biological decision-making scenarios. Thus, connecting the geometry of cell-fate decisions to their underlying regulatory networks remains an open question. Recently, two regulatory networks have been shown to exhibit tristability – a toggle switch with self-activation, and a toggle triad. Here, we show that these two motifs are distinct from a geometrical point of view, and identify two bifurcations in their behaviour: the standard cusp and the elliptic umbillic. We study experimentally accessible signatures of the differences in tristability between the two motifs. We also show how the standard cusp can be used to quantitatively model cell fate transition data for the Epithelial to Mesenchymal transition on TGF-*β* induction in the context of cancer cells. Our work uncovers geometrical signatures of gene regulatory motifs and demonstrates how different gene regulatory networks can encode tristability in dynamically distinct ways.

## 1 Introduction

Recent theoretical work has formalized the idea of Waddington’s landscape and classified the possible properties of the dynamical behaviour of systems with three stable states [1]. This classification is quite general and independent of the details of the underlying differential equations. The utility of this approach has been demonstrated in the context of cell fate decisions. In this geometric approach, the coordinates are abstract, cell fates correspond to stable fixed points or attractors and decisions correspond to saddle points [2]. The associated unstable manifolds demarcate the stable regions of different fates. For example, the cell fate decision where a cell originally in a central attractor can differentiate into two different possible fates has been classified into two distinct possibilities: the dual cusp and the heteroclinic flip. The difference between these two decisions has now been studied in various contexts [3, 4, 5, 6, 7].

However, much of this work has directly tried to infer geometric models from data on cell fates and the connection of these landscapes to underlying gene regulatory networks is not yet clear. While models of gene regulatory networks suffer from known problems of over-parameterization, they have the advantage of directly parameterizing genes important to the decision and thus continue to be a popular modeling choice. Hence, it is important to study whether different kinds of gene regulatory networks models can implement the dynamical possibilities predicted earlier [1]. Conversely, a geometric approach can be helpful to clarify the differences in tristability between different regulatory networks. Further, it can encourage more quantitative experiments in the corresponding biological systems where the geometrical model is well-suited to quantitatively fit cell fate data.

Gene regulatory motifs have been commonly used to model cell fate decisions in various contexts including development, cancer and microbial behaviour [8, 9, 10, 11]. The toggle switch, initially proposed as a synthetic circuit in *E. coli* [12], which involves the mutual repression of two elements, is a motif that has appeared in diverse contexts, with the two elements mostly representing transcription factors. The transcription factors PU.1 and GATA1 are believed to constitute a toggle switch and repress each other’s expression in the context of hematopoietic cell lineage decisions [13, 14, 15]. The transcription factors NANOG and GATA6 are similarly believed to constitute a mutually repressive circuit in early mouse development [16, 17, 18]. Typically, the toggle switch has two stable states, with one of the two genes being highly expressed in the stable state. Thus, a toggle switch between two elements A and B often have (high A, low B) and (low A, high B) as the two stable states.

In some cases, the toggle switch can have elements which are self-activating. This motif - a Toggle Switch with Self Activation (TSSA) - can allow for a third stable state which has intermediate levels of both the elements - (medium A, medium B) [19]. For instance, in the context of the Epithelial to Mesenchymal Transition (EMT) in cancer cells, the core regulatory circuit of EMT involves a micro-RNA miR-200 and a transcription factor ZEB1, which is self-activating [20]. This circuit predicts the presence of a hybrid state which is intermediate between the epithelial and mesenchymal states. The existence of this hybrid state was first predicted theoretically and has since been experimentally confirmed [20, 21, 22]. The self-activating terms in the toggle switch are crucial to obtaining tristability [23].

Subsequently, attention has gone to mutually repressive circuits with a larger number of components including the toggle triad (TT) which involves three mutually repressive components, a circuit important in the context of CD4 expressing T cells which differentiate into multiple lineages [24, 25, 26]. The toggle triad also has tristability, where each stable state typically involves the high expression of one component and the down-regulation of the other two. For a toggle triad with three components A, B, C, the stable states are typically (high A, low B, low C), (low A, high B, low C) and (low A, low B, high C). The study of such tristable gene regulatory networks continues to be an active area of research [11, 27, 18].

Here, we study the dynamics of these gene regulatory motifs using a geometrical approach and study tristability in the context of the TSSA and the TT. We identify the potential landscapes which have the same decision structure as these motifs and show that they correspond to two distinct geometries. We also examine the bifurcation diagrams that result from varying different parameters in the motif and relate them to the universal bifurcation diagrams obtained on varying two parameters predicted earlier. We show how the geometric picture allows us to suggest potential experiments that can distinguish between the behaviour of the two motifs. We also show how available experiments in the case of EMT, believed to be governed by the TSSA, can be fit using the corresponding geometrical model. Finally, we make predictions for systems whose master regulators have a TT motif.

## 2 Results

### 2.1 The TSSA and TT have distinct landscapes

The two gene regulatory motifs we study here, TSSA and TT, are both known to have three stable phenotypic states. These correspond to stable fixed points in the flow. We find that the flow of the two motifs has two distinct landscapes. The first (corresponding to TSSA), shown in Fig 1 A (i), is a modified form of the so called butterfly potential. There is a flow corresponding to the landscape shown in Fig 1 A (ii). As the TSSA motif has only two state variables, its flow can be directly plotted as shown in Fig 1 A (iii). There are three stable states corresponding to (high A, low B), (low A, high B), and (medium A, medium B). Even though the flow is in two dimensions, the geometry is effectively one dimensional as there is a single curve connecting all the five fixed points. This structure has important consequences for the phenotypic behaviour of the TSSA as we show later. The modified butterfly potential we use also has three stable fixed points with two saddle points in between and is in fact one-dimensional though it is plotted in two dimensions here for visualization purposes. Therefore, the geometric structure of the butterfly potential is qualitatively the same as that of the TSSA.

**Figure 1.**
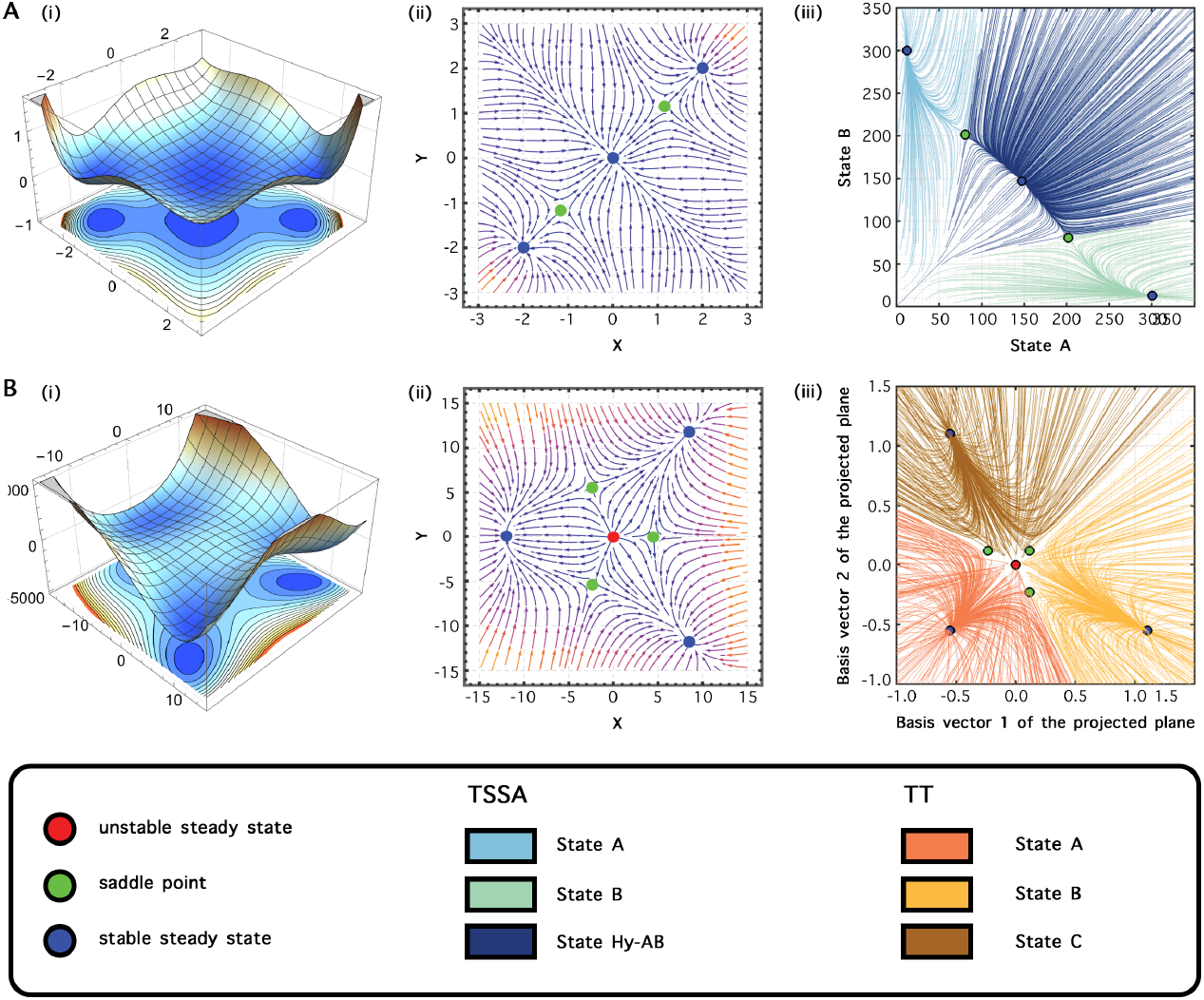
Geometric models analogous to TSSA and TT. (A)(i) Potential landscape for the modified butterfly equation (Methods). (ii) Flow diagram for the modified butterfy equation. (iii) Flow diagram for TSSA. (B)(i) Potential landscape for the elliptic umbilic potential. (ii) Flow diagram for the elliptic umbilic potential. (iii) 2D projection of flow diagram for TT.

The dynamics of the TT motif is described by a different geometry given by modifying the Elliptic Umbilic as shown in Fig 1 B (i). The flow corresponding to the landscape is plotted in Fig 1 B (ii). Since the TT motif has three state variables, its flow cannot be directly plotted in 2 dimensions. However, we project the dynamics of the TT onto the plane formed by the three saddle points shown in green in Fig 1 B (ii). Here,the three states correspond to (high A, low B, low C), (low A, high B, low C) and (low A, low B, high C) but the coordinates are a linear combination of the three genes. The potential now is intrinsically two-dimensional as there is no way to represent it in one dimension. It has three stable fixed points and three saddle points with an unstable fixed point in between. This is qualitatively similar to the ellipitic umbilic which has the same set of fixed points and stability properties.

While both the landscapes have three stable fixed points, the connections between the fixed points are clearly distinct in the two cases. Since the TSSA is effectively one dimensional, there is no direct connection between the two farthest fixed points. In contrast, all of the fixed points in the landscape associated with TT are connected to each other with a saddle point.

This difference has important consequences for the kinds of bifurcation diagrams obtained by varying parameters on these landscapes. In 2-dimensional parameter space, it is known that lines of saddle-node bifurcations can either meet in a cusp or intersect transversely. This allows a classification of the possible bifurcation diagrams obtained by varying 2-parameters. In Fig 2 we show the bifurcation diagrams obtained by varying different combinations of 2 parameters in the dynamics of the TSSA and the TT motif. We find that the particular bifurcation diagram obtained depends on the choice of parameters. Here, we show a couple of illustrative examples of these bifurcation diagrams. For the TSSA, motivated by experimental observations that vary fate by varying an external signal, we show the two dimensional bifurcation diagrams with the external signal and threshold of self-activation of A as well as the fold-change of self-activation of A. In the first case, we obtain a *dual cusp*, a point where two saddle points and a stable fixed point merge into a saddle. In the second case we obtain a *standard cusp*, a point where two stable fixed points and a saddle point merge into a stable fixed point. While the nature of the two dimensional diagrams is different, the regions traversed when going along the horizontal axis (corresponding to increasing the external signal) can be the same depending on the value of the threshold or fold-change.

**Figure 2.**
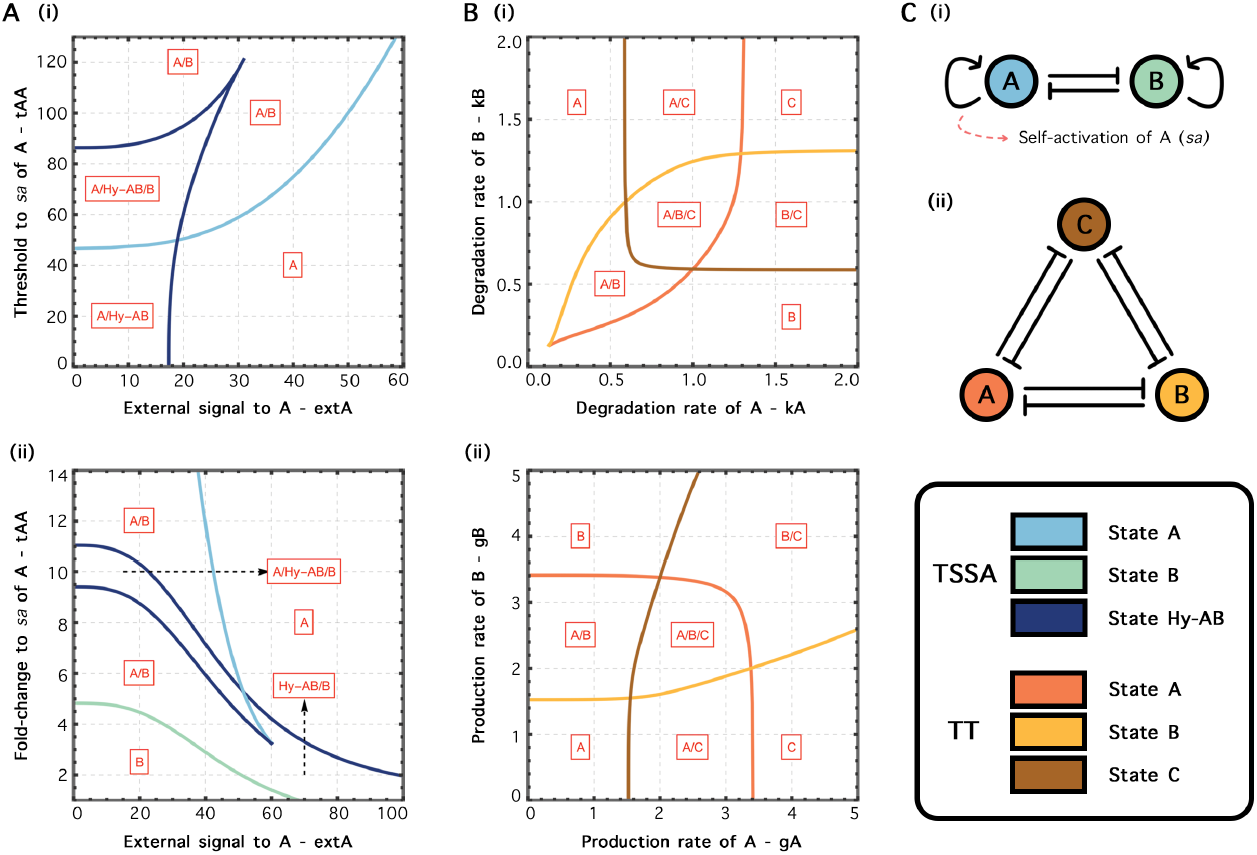
Bifurcation diagrams for TSSA and TT. (A)(i) A two dimensional bifurcation diagram with the threshold parameter of self-activation and the external signal activating A chosen as parameters. (ii) A bifurcation diagram with the fold-change parameter of self-activation of A and the external signal activating A chosen as parameters. (B)(i) Bifurcation diagram with the production rate of A and the production rate of B chosen as parameters. (ii) Bifurcation diagram with the degradation rate of A and the degradation rate of B chosen as parameters. (C)(i) Schematic for the network of TSSA. (ii) Schematic for the network of TT.

In the case of the TT, the bifurcation diagram is given by the elliptic umbilic. All lines of saddle node bifurcations intersect transversely with each other. The tristable region is flanked by three bistable regions, which in turn have a monostable region in between them. All monostable regions are connected to each other with an intervening bistable region as shown in Fig. 2 B.

### 2.2 Signaling perturbations lead to distinct trajectories of phenotypes

The distinct landscapes and bifurcation diagrams obtained for the TSSA and TT lead to very different phenotypic trajectories when one of the nodes is activated by an external signal. Since the TSSA is essentially one dimensional, trajectories going from (low A, high B) to (high A, low B) on activation of A have to necessarily go through the intermediate ((medium A, medium B)) state. As shown in Fig. 3, a population of cells starting at (low A, high B) that are pushed towards (high A, low B) by a signal, travel through the intermediate or hybrid state. Importantly, independent of the details of signal, any trajectory from (low A, high B) to (high A, low B) must go through the region which constitutes the domain of attraction of the (medium A, medium B) state.

**Figure 3.**
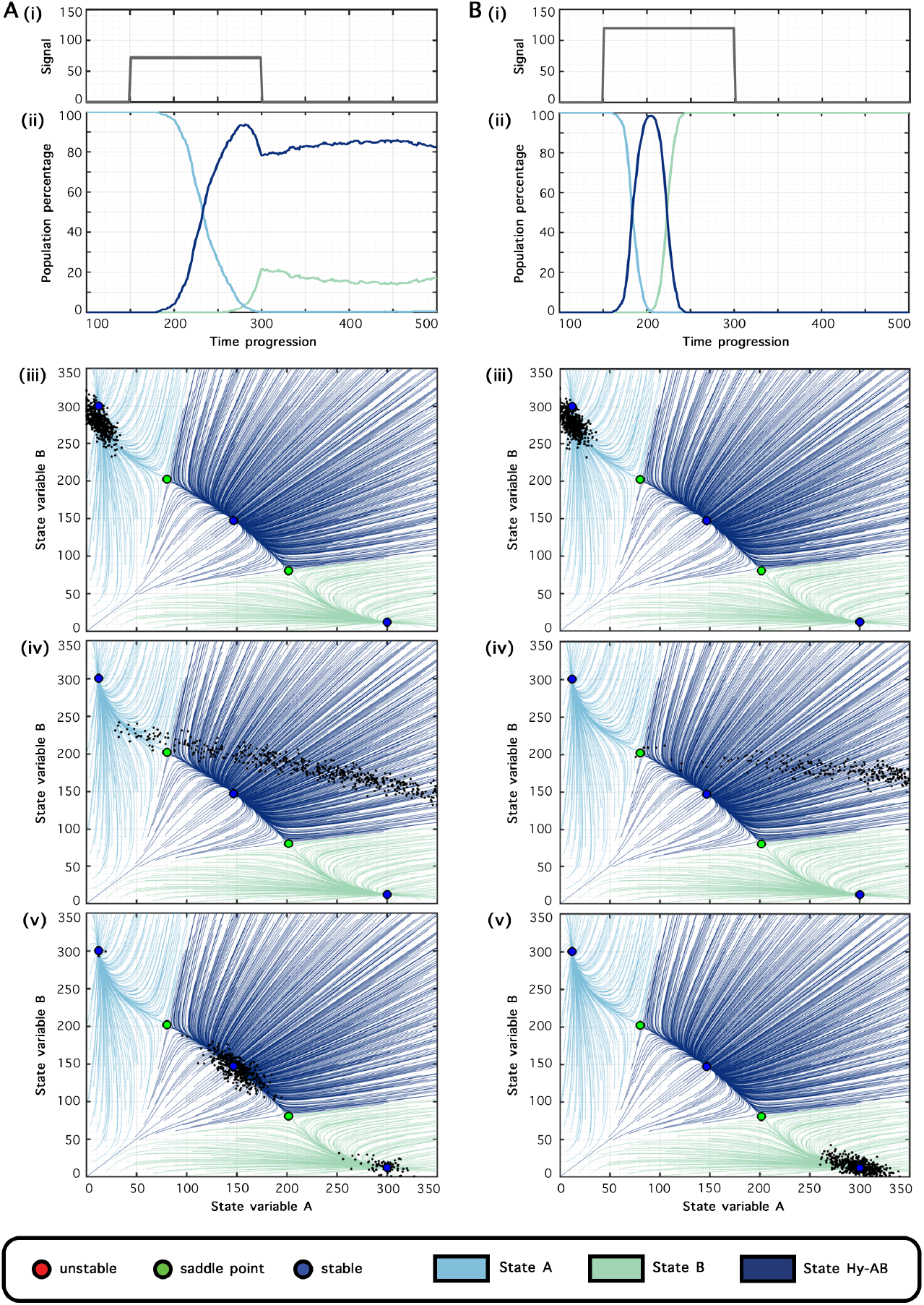
Population redistribution by an external signal - TSSA. (A)(i) External activation provided to gene B as a rectangular signal. (ii) Change in population percentages of the states as a function of time. (iii) Snapshot of the population (black dots) overlaid on the flow diagram before the rectangular signal. (iv) Snapshot of the population during the rectangular signal. (v) Snapshot of the population after the rectangular signal. (B) Same as (A) for a rectangular signal of higher strength (*as seen in A (i) vs B (i)*).

Here, we show the population percentages, as classified by the domain of attraction in the absence of the signal, for two different signal strengths in Fig. 3. We also plot three snapshots of the phase space for a group of cells in Fig. 3 A and B for two different signal strengths. The frequency of (low A, high B) state first decreases, then the hybrid (medium A, medium B) population increases and finally the (high A, low B) population increases. The speed of cell fate determination and the population percentages can vary depending on specific signal but the ordering is fixed by the geometry. For example, there is no combination of activation of A and repression of B that can reverse the order of appearance of the (medium A, medium B) and the (high A, low B) state.

The situation is very different in the case of the TT. Here, the order of appearance as well as the percentage of populations depend on the precise nature of the signal. Hence, starting from the (high A, low B, low C) state and activating both B and C leads to the near simultaneous appearance of the (low A, high B, low C) and (low A, low B, high C) states as shown in Fig 4 A. On the other hand, a signal which activates only the B state leads only to the appearance of the (low A, high B, low C) and the (low A, low B, high C) state is never occupied by the cells. We can see this clearly by projecting the flow on to two-dimensions as before. The trajectories taken by a group of cells in the two cases is shown in Fig 4 A (iii)-(v) and Fig 4 B (iii)-(v).

**Figure 4.**
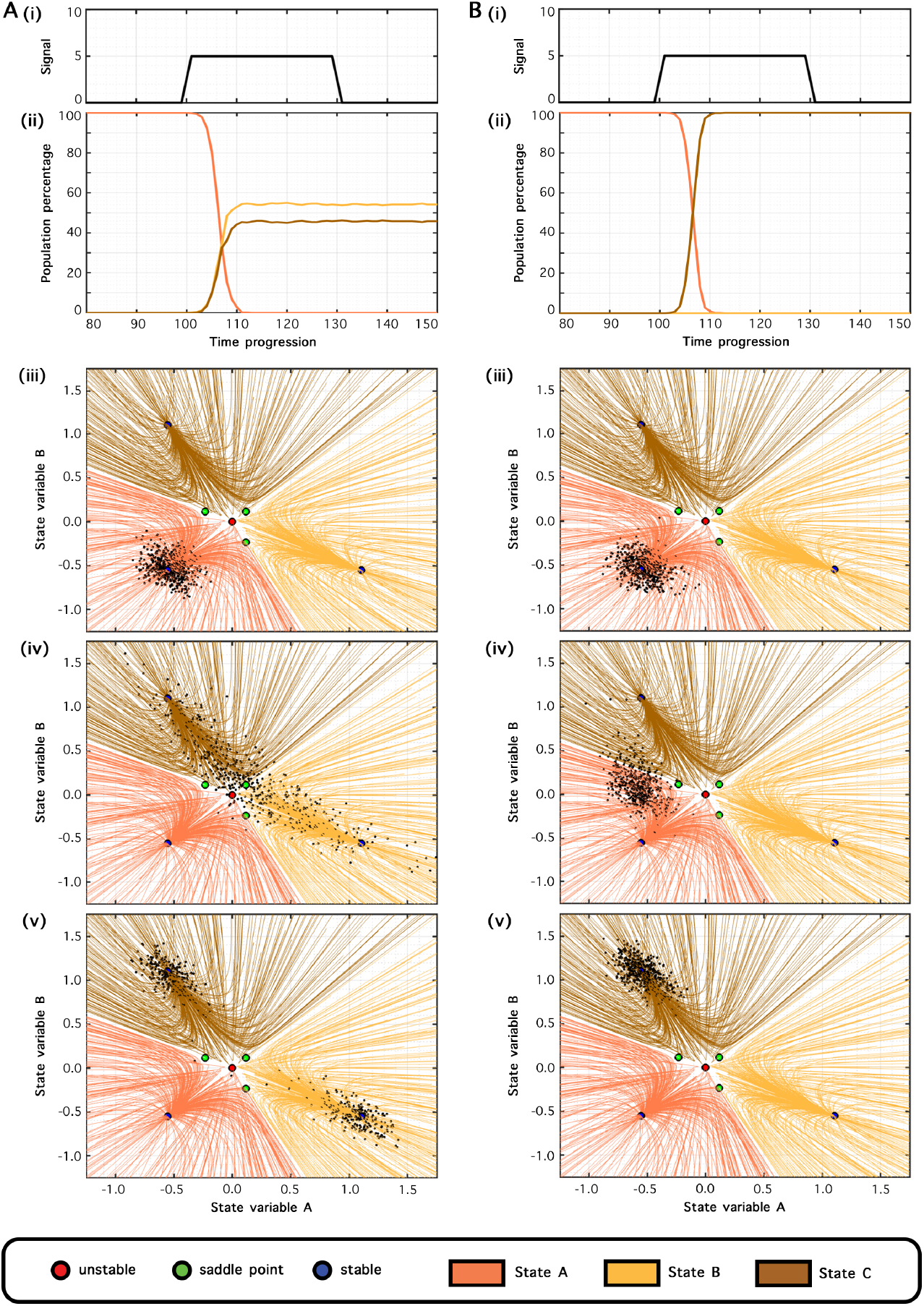
Population redistribution by an external signal - TT. (A)(i) External activation provided to gene B and C as a rectangular signal. (ii) Change in population percentages of the states as a function of time. (iii) Snapshot of the population (black dots) overlaid on the flow diagram before the rectangular signal. (iv) Snapshot of the population during the rectangular signal. (v) Snapshot of the population after the rectangular signal. (B) Same as (A) for a rectangular signal activating only C.

**Figure 5.**
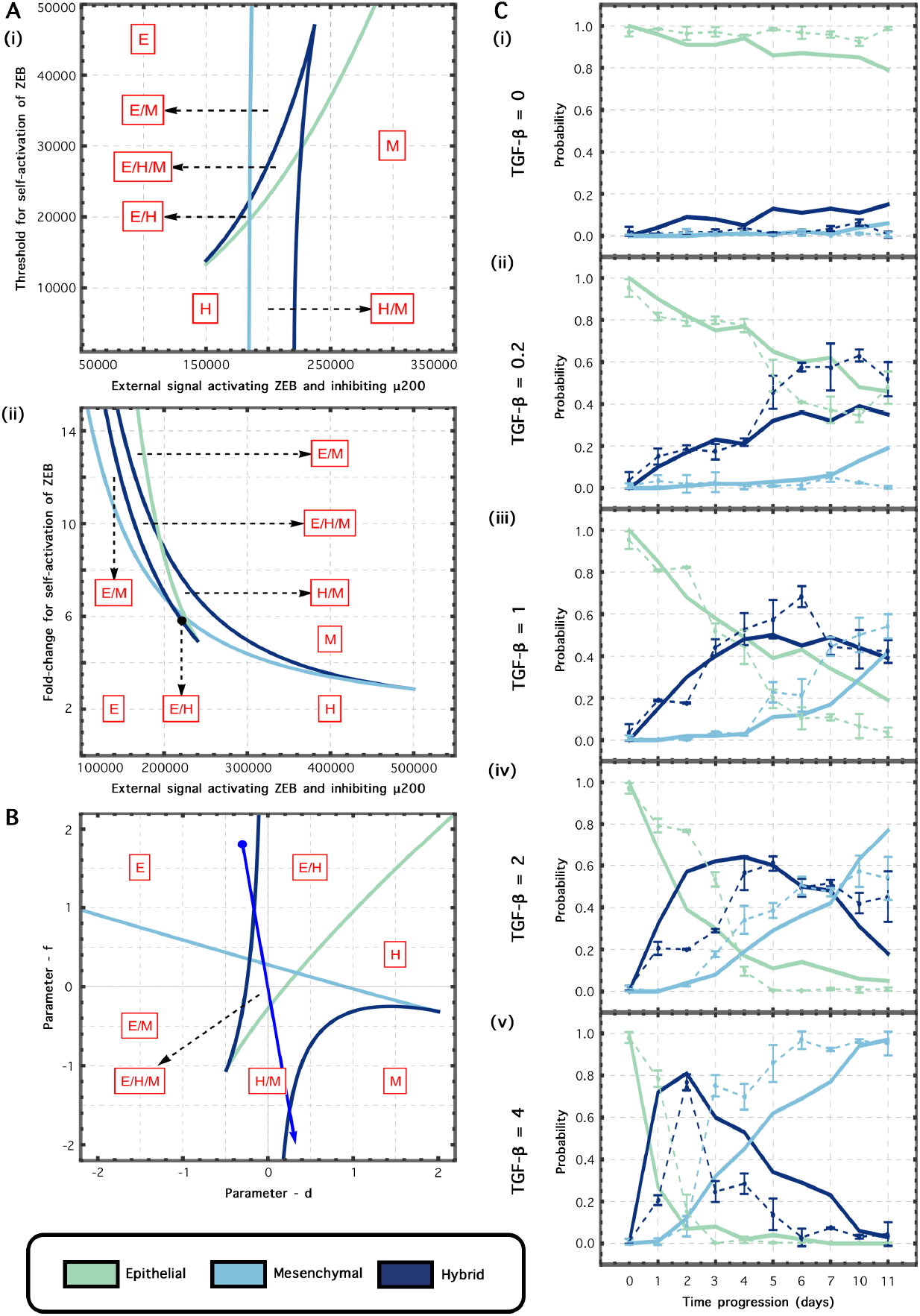
Fitting parameters of geometric model to EMT data. (A)(i) Bifurcation diagram with the threshold parameter of self-activation of ZEB and the external signal activating ZEB and inhibiting *µ*200. (ii) Bifurcation diagram with the fold-change parameter of self-activation of ZEB and the external signal activating ZEB and inhibiting *µ*200. (B) The bifurcation diagram obtained by varying two parameters - *d* and *f* in the butterfly equation is similar to the one obtained in the gene regulatory network of EMT dynamics. The blue path shown represents the path followed for the fitted parameters to data. (C)(i)-(v) Data corresponding to progression of population percentages of Epithelial, Mesenchymal and Hybrid populations at ten time points for different concentrations of TGF-*β* (shown in dashed line). The data is from Ref. [28]. The population percentages as determined at the best fit (shown in solid line).

The observation of the phenotypic distribution in response to a signal gives a clear experimentally testable distinction between tristability for these two kinds of motifs. A population curve of the form of Fig. 4 A (ii) for example suggests that it is unlikely that a one-dimensional landscape governs the system.

### 2.3 Cell fate data in EMT can be quantitatively described by a landscape

As mentioned previously, EMT in cancer cells is believed to be governed by a TSSA. The appearance of the one-dimensional landscape in the TSSA suggests that cell fate transitions in EMT can be modeled geometrically using a butterfly potential. To first verify this, we plotted the bifurcation diagrams corresponding to a model of the core regulatory circuit of EMT taken from Ref. [29]. The core circuit is composed of miR-200 and ZEB whose regulatory structure is similar to the TSSA. An exogenous signal of TGF-*β* likely activates SNAIL which in turn activates ZEB, whose activation promotes the mesenchymal fate. This TSSA can allow for three states - epithelial (high miR-200, low ZEB1), mesenchymal (low miR-200, high ZEB1) and hybrid epithelial/mesenchymal (medium miR-200, medium ZEB1). The bifurcation diagram is shown in Figure 6A with the external signal taken to activate ZEB and inhibit miR-200 (which SNAIL does). As before, the nature of the bifurcation diagram depends on the particular parameters chosen but we notice a standard cusp governing the transition in both cases.

**Figure 6.**
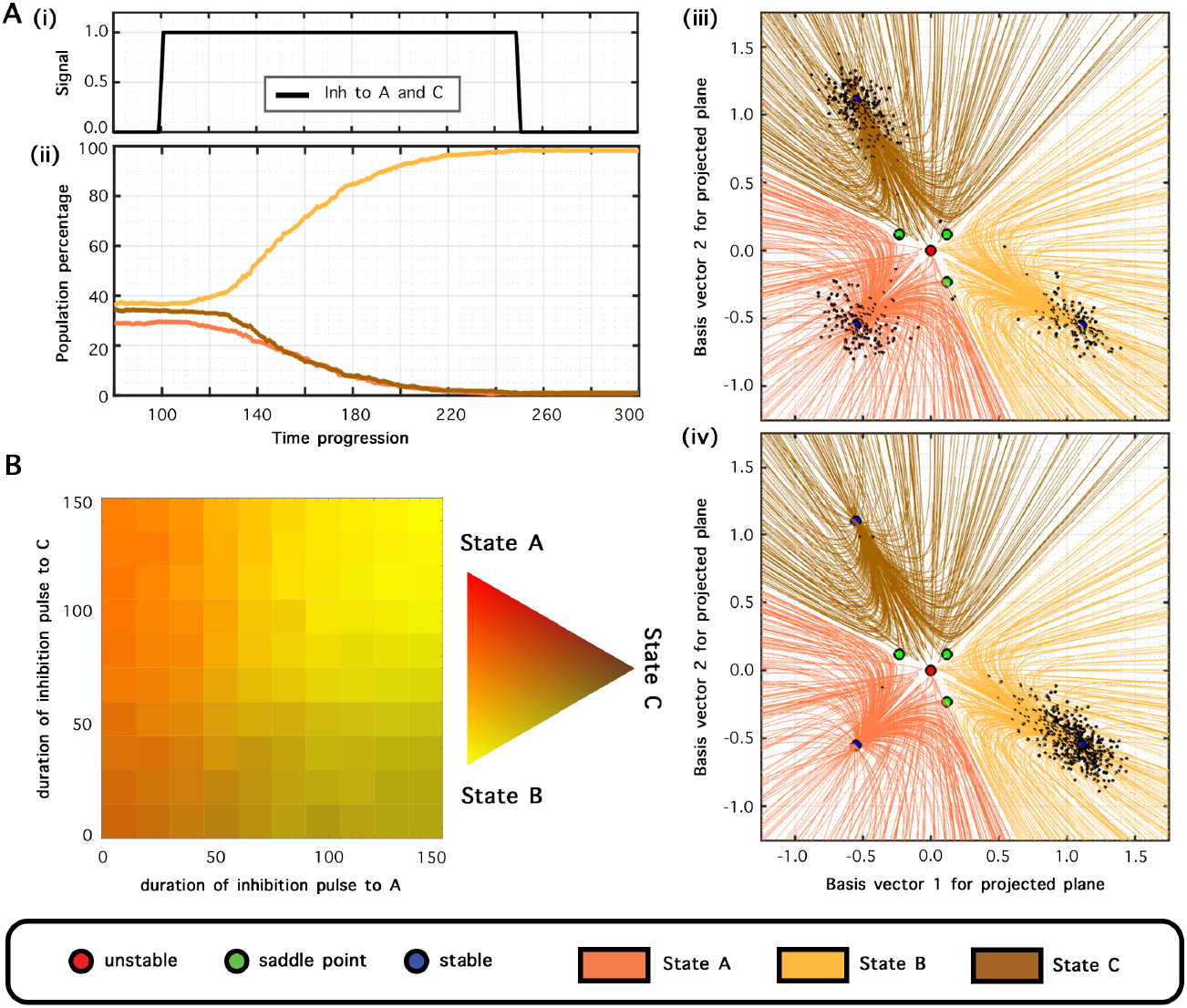
Effects of time duration of external inhibition on genes A and C. (A)(i) External activation provided to gene B and inhibitions provided to genes A and C as different rectangular signals. (ii) Change in population percentages of the states as a function of time. (iii) Snapshot of the population (black dots) overlaid on the flow diagram before the rectangular signal. (iv) Snapshot of the population after the rectangular signal. (B) Heatmap indicating the population distribution for different pairs of values for the duration of external inhibition to genes A and C.

Ref. [28] investigated the transition from the Epithelial to Mesenchymal state on TGF-*β* induction and quantitatively characterized the cell fates. They observed the dynamics of appearance of the Epithelial, Hybrid and Mesenchymal states for different concentrations of TGF-*β* using time-course flow cytometry. They found a sequential appearance of the Hybrid state followed by the Mesenchymal state, which is consistent with a one-dimensional potential landscape. Both of these observations suggest that it should be possible to model this decision using a one-dimensional potential.

We model the transition using the aforementioned butterfly potential in one dimension (Methods). On varying two of the parameters of the potential, we get a bifurcation diagram which also has a standard cusp as shown in Figure 5B. Two parameters of the potential are taken to be linearly dependent on the TGF-*β* concentration, as has been done in previous studies in other contexts. The potential has tristability and the three stable states are taken to correspond to the Epithelial, Hybrid and Mesenchymal phenotypes. Gaussian noise is added to the model to simulate stochasticity in gene regulation.

The dependence of the coefficients on the TGF-*β* concentration and the strength of the noise constitutes the parameters to be fit in the model. We find that we are able to capture the cell fate data across concentrations of TGF-*β* with our simple model (Figure 5C). The fit implies that the parameters change as the TFG-*β* concentration changes and this dependence is shown in the solid blue line in Fig 5B. The parameters are initially in the monostable region corresponding to the Epithelial fate only (for no TGF-*β*). They then pass through the bistable region corresponding to Epithelial and Hybrid states, then the tristable region containing all 3 states, followed by bistable region comprising hybrid and Mesenchymal states, followed by the monostable Mesenchymal region at increasing concentrations of TGF-*β*. Similar paths are possible in the bifurcation diagrams for the gene regulatory model of ZEB and miR-200 (Supplementary Information). There is a range of parameter values which fit the data equally well. Another set of parameters along with the phenotypic behaviour is shown in SI Figure 2.

### 2.4 Transitions in the Toggle Triad require two inhibitory signals

For the Toggle Triad, our landscape analogy helps us confirm qualitative results and make quantitative predictions. Because of the nature of the landscape and the inherent two-dimensional nature of the geometry, precise control over fate requires two signals. A single signal is likely to lead to mixed states in the Toggle Triad as we demonstrate below.

In Fig. 6, we demonstrate the effect of varying the duration of inhibition to state A and C starting from an initial condition which is in a mixture of the three states. Fig. 6A shows the transition from all three states being populated to only B being populated. If we inhibit only one of the states, we typically get mixed states, so inhibition of only A leads to states mixed between (low A, low B, high C) and (low A, high B, high C) whereas inhibition of C leads to mixed states of (high A, low B, low C) and (low A, high B, low C). Only on inhibition of both A and C do we get all cells to be in state (low A, high B, low C) as shown in Figure 6B.

The fact that two signals are needed to control a state is reflective of the difference between the TT and TSSA. In the switch, since the geometry is essentially one dimensional, any signal essentially gets projected on to that one dimension. In the triad, there are fundamentally two degrees of freedom and two signals can be used to control the motion in a two-dimensional landscape. This is consistent with qualitative observations that the differentiation of CD4+ T cells into a particular state require treatment with multiples cytokines [30] suggesting that the elliptic umbilic may be the right geometry to fit cell fate data for this system.

## 3 Discussion

Here, we have shown that a landscape picture of the two gene regulatory motifs allows distinguishing the essential difference between their tristability. We find that the TSSA can be modeled with a one-dimensional potential whereas the TT requires a two-dimensional potential. All possible transitions between cell fates are allowed in the TT but not in the TSSA. This fact can be seen experimentally with perturbation experiments as we demonstrate *in silico* consistent with previous suggestions [31, 4]. Hence, the decision structures of the two motifs are fundamentally different and empirically distinguishable. Nevertheless, we find that the particular bifurcation diagram is not directly associated with the topology of the motif and depends on the parameters being varied. For example, we find both the dual cusp and the standard cusp in the TSSA depending on the parameters that are varied.

The stability of a hybrid phenotype in EMT was confirmed, in part, as a result of mathematical modeling [20, 32]. However, quantitative data on cell fate in EMT has not been fit before partly because gene-regulatory models are over-parameterized for such quantitative fits. Here, we show that the geometry of EMT is governed by a standard cusp. We then fit a simple one-dimensional potential landscape with a standard cusp to cell fate data in the presence of exogenous TGF-*β*. We expect the standard cusp to be important in cases where the topology is one-dimensional and the decision moves from one attractor to another through a hybrid state.

The cusp is a special point in parameter space. Going around the cusp would allow the transition between cell states (e.g. from Epithelial to Hybrid) without an intervening bifurcation. It remains to be seen in future work whether such transitions are biologically plausible and our fit to EMT data does not explore this region.

We show that the Toggle Triad is governed by a different topology called the elliptic umbilic. The elliptic umbilic allows all possible binary transitions between the three states. We expect it to be important in cases where a progenitor state can differentiate into one of three types. Our framework also makes predictions for experiments that perturb fates in such a system. We predict that it should be possible to obtain different sequence of phenotypes going from one state to the other two depending on the strength of the inhibition to two of the states. For CD4+ T cell differentiation where the master regulators are believed to have a toggle triad topology, it is possible to implement this inhibition using cytokines [33]. Our landscape should be able to quantitatively fit data on such inhibition experiments and predict novel outcomes.

There have been recent efforts to fit geometric landscapes to single cell data [5, 34]. It would be interesting to study if the landscapes we identify for these two different gene regulatory motifs can be identified in such high-dimensional data for the corresponding biological systems.

The small gene regulatory networks we consider have received extensive study and we show that they correspond to distinct geometries predicted in Ref. [1]. However, these networks are only approximations to much more complex regulatory networks. Recent work suggests that they can be understood as coarse-grained approximations to larger networks which nevertheless correctly characterize the low-dimensional phenotypic behaviour at the level of cell fate [35, 36]. The geometry does not depend on interpretation of the variables in underlying differential equations. Hence, even if the equations describe effective interactions between “teams” or “modules”, the landscapes we identify will still govern the transitions. Geometric models describe the low-dimensional phenotypic behaviour most parsimoniously. We have shown here that even for small networks, it is possible to reduce the dimension and parameters required to understand the decision. It remains an open question if these landscapes can be understood to emerge from higher dimensional regulatory networks.

## 4 Methods

### Equations for potential landscapes

The mathematical form for the butterfly and elliptic umbilic used are given as follows. For the butterfly:

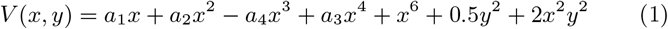

The butterfly equation, which typically has only one variable (x) is modified here to include two variables (x and y). For the elliptic umbilic:

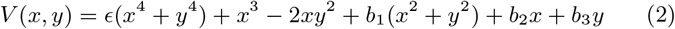

A small quartic term is added to keep the potential bounded. The flow lines and potential landscapes in Fig 1A and 1B are drawn using the above equations. Parameter values used are a_1_ = 0, a_2_ = 1, a_3_ = −2, a_4_ = 0 and ϵ = 0.1, b_1_ = −10.8, b_2_ = −0.8 and b_3_ = −2.4.

### Ordinary Differential Equation model

The interaction between two genes in the network given by *A* and *B* is modeled using the following generalized Hill functions

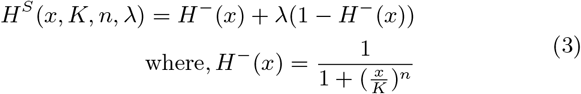

The generalized Hill function has three parameters.*K* is the threshold parameter, *n* is the corresponding Hill coefficient, and *λ* is the maximum fold-change. Values of *λ <* 1 imply inhibition and of *λ >* 1 imply activation. Therefore, both inhibition and activation can be modeled using the same function by changing the value of *λ*.

The equations for the TSSA network are as follows:

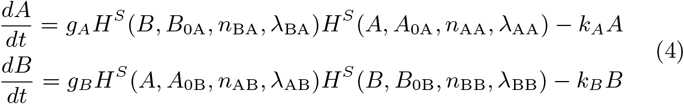

where *g*_*A*_ and *g*_*B*_ are basal production rates and *k*_*A*_ and *k*_*B*_ are degradation rates. The equations for the TT network are the following:

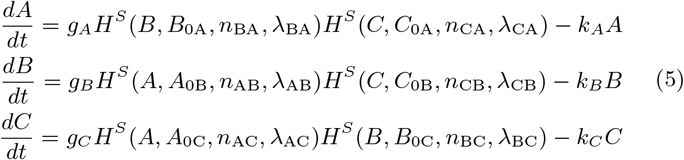

The equations used to create the bifurcation diagrams in Figure 5 are similar but more complex. They are taken from Ref. [29]. In particular, we use Equations 1_1_, 2 and 3 in the Supplementary Information of the above reference with parameter values taken from Table S1.

### Flow and Bifurcation diagrams

Since the Toggle Triad has a three-dimensional state space, we draw the flow lines in two dimensions as in Figure 1B(iii) by projecting the flow onto the plane formed by the three saddles. We solve Equation 5 for different initial conditions, and project the three-dimensional trajectory obtained on to the plane formed by the three saddles. Trajectories are coloured by the fixed point that they end up at.

All two dimensional bifurcation diagrams as in Figure 2 are obtained by solving two sets of algebraic equations. The first equation is obtained by setting the left hand side of Equation 4 or Equation 5 equal to zero. The second is obtained by requiring that there is at least one zero eigen-value in the system for which we calculate the determinant of the Jacobian of Equation 4 or Equation 5 respectively and set it equal to zero. These equations are solved by holding one parameter fixed and numerically solving for the other. For example, in Figure 2B(i), we can solve for *k*_*B*_ while keeping the value of *k*_*A*_ fixed. As *k*_*A*_ is varied, this leads to a line of saddle node bifurcations.

### Modeling External Signal

We model the external signal using the Hill function. For example, for the TSSA, the modified equations are

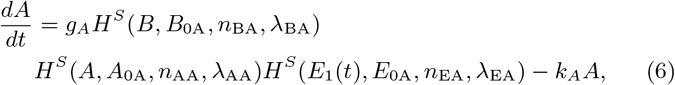

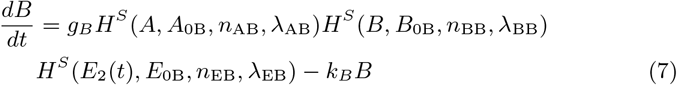

where *E*_1_(*t*) and *E*_2_(*t*) are time-dependent signals which can be activatory or inhibitory depending on the value of the threshold parameter *λ* in the Hill function. The time dependence is modeled using a rectangular function which has value *S* between two time points *T*_1_ and *T*_2_ and is zero otherwise. External signals are modeled in an identical fashion for the TT. Parameter values are provided in the Supplementary Information.

### Classification of Fates

To classify fates in the TSSA, we find the eigenvectors near the two saddle points. We then extend the stable direction to the saddle linearly in both directions. This partitions the phase space into four regions. We associate the regions containing the (high A, low B), (low A, high B) and hybrid fixed point (medium A, medium B) with the corresponding fates. A fourth region corresponding to low expression values in both A and B is irrelevant in practice. This allows us to approximate the stability domains and classify fates.

For the TT, we simply use the symmetry of our system to associate the fate with (high A, low B, low C), (low A, high B, low C) or (low A, low B, high C) depending on which expression value is the highest.

### Fitting the data

For Fig. 5C, the data is taken from Ref. [28] Fig 6C. TGF-*β* values in the original data are in ng/ml and time is in days.

To fit the data, we use the potential given below

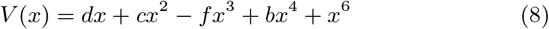

We set *c* = 1 + *d* and *b* = −2 giving us only two parameters *d* and *f* to vary. Their dependence of TGF-*β* is modeled as

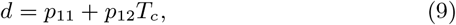

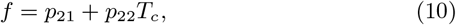

where *p*_11_, *p*_12_, *p*_21_ and *p*_22_ are parameters to be fit and *T*_*c*_ is the TGF-*β* concentration. The initial condition is sampled using a normal distribution with mean *x*_0_ set to the position of the first attractor (corresponding to the E state) and standard deviation 0.2.

We then solve the equations

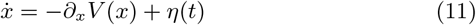

where *η*(*t*) is a Gaussian white noise given by ⟨*η*(*t*)*η*(*t′*) = *gδ*(*t* − *t′*).

The fates are decided by a threshold if *x > θ* or *x <* −*θ* with *θ* = 0.5 corresponding to the Epithelial and Mesenchymal states with intermediate states classified as hybrid. We use a time-step *dt* = 0.01 and time is rescaled arbitrarily so that the total time is *T* = *τ/*(15 ∗ *dt*) with *τ* = 1.

In principle, we have 5 remaining parameters to fit but we find that *p*_12_, *p*_22_ and *g* are the most important ones. We find *p*_11_ = −0.2 and *p*_21_ = 0.32, *p*_12_ = 1.2, *p*_22_ = −1.95, and *D* = 1.59 to fit well by uniformly sampling the parameter space. Since the number of parameters are small and meaningful values of the parameters are confined to a relatively small region, such a uniform sampling is not very computationally expensive.

## Acknowledgements

AR acknowledges support from the Department of Atomic Energy, Government of India (under project RTI4006) and the Simons Foundation (287975). MKJ was supported by Param Hansa Philanthropies. ASD acknowledges support by Prime Ministers’ Research Fellowship (PMRF) awarded by the Government of India.

