## Supplementary Information for "Distinct geometrical landscapes distinguish between modes of tristability in gene regulatory networks"

### Distinct modes of tristability in gene regulatory motifs: Supplementary Information

October 11, 2025

#### 1 Parameter Values used for TSSA and TT

Table 1 and Table 2 give the parameter values used for the TSSA and TT throughout the text. For Figure 3 in the Main text, we use the following parameter values for the external signal $E_{0\mathrm{A}} = 100$, $\lambda_{\mathrm{EA}} = 10$, $n_{\mathrm{EA}} = 3$, $E_{0\mathrm{B}} = 100$, $\lambda_{\mathrm{EB}} = 0.2$, $n_{\mathrm{EB}} = 3$. For Figure 5 in the main text, we use the following values $E_{0\mathrm{A}} = 1$, $\lambda_{\mathrm{EA}} = 0.4$, $n_{\mathrm{EA}} = 4$, $E_{0\mathrm{C}} = 1$, $\lambda_{\mathrm{EC}} = 0.4$, $n_{\mathrm{EC}} = 4$.

Table 1: Parameter Values for Toggle Triad

| Parameter | Value |
|---|---|
| $g_{\mathrm{A}}$ | 2 |
| $B_{0\mathrm{A}}$ | 1 |
| $n_{\mathrm{BA}}$ | 4 |
| $\lambda_{\mathrm{BA}}$ | 0.1 |
| $C_{0\mathrm{A}}$ | 1 |
| $n_{\mathrm{CA}}$ | 4 |
| $\lambda_{\mathrm{CA}}$ | 0.1 |
| $k_{\mathrm{A}}$ | 1 |
| $g_{\mathrm{B}}$ | 2 |
| $A_{0\mathrm{B}}$ | 1 |
| $n_{\mathrm{AB}}$ | 4 |
| $\lambda_{\mathrm{AB}}$ | 0.1 |
| $C_{0\mathrm{B}}$ | 1 |
| $n_{\mathrm{CB}}$ | 4 |
| $\lambda_{\mathrm{CB}}$ | 0.1 |
| $k_B$ | 1 |
| $g_{\mathrm{C}}$ | 2 |
| $A_{0\mathrm{C}}$ | 1 |
| $n_{\mathrm{AC}}$ | 4 |
| $\lambda_{\mathrm{AC}}$ | 0.1 |
| $B_{0\mathrm{C}}$ | 1 |
| $n_{\mathrm{BC}}$ | 4 |
| $\lambda_{\mathrm{BC}}$ | 0.1 |
| $k_B$ | 1 |

Table 2: Parameter Values for Toggle Switch with Self Activation

| Parameter | Value |
|---|---:|
| $g_{\mathrm{A}}$ | 5 |
| $B_{0\mathrm{A}}$ | 120 |
| $n_{\mathrm{BA}}$ | 1 |
| $\lambda_{\mathrm{BA}}$ | 0.1 |
| $A_{0\mathrm{A}}$ | 80 |
| $n_{\mathrm{AA}}$ | 3 |
| $\lambda_{\mathrm{AA}}$ | 10 |
| $k_{\mathrm{A}}$ | 0.15 |
| $g_{\mathrm{B}}$ | 5 |
| $A_{0\mathrm{B}}$ | 120 |
| $n_{\mathrm{AB}}$ | 1 |
| $\lambda_{\mathrm{AB}}$ | 0.1 |
| $B_{0\mathrm{B}}$ | 80 |
| $n_{\mathrm{BB}}$ | 3 |
| $\lambda_{\mathrm{BB}}$ | 10 |
| $k_B$ | 0.15 |

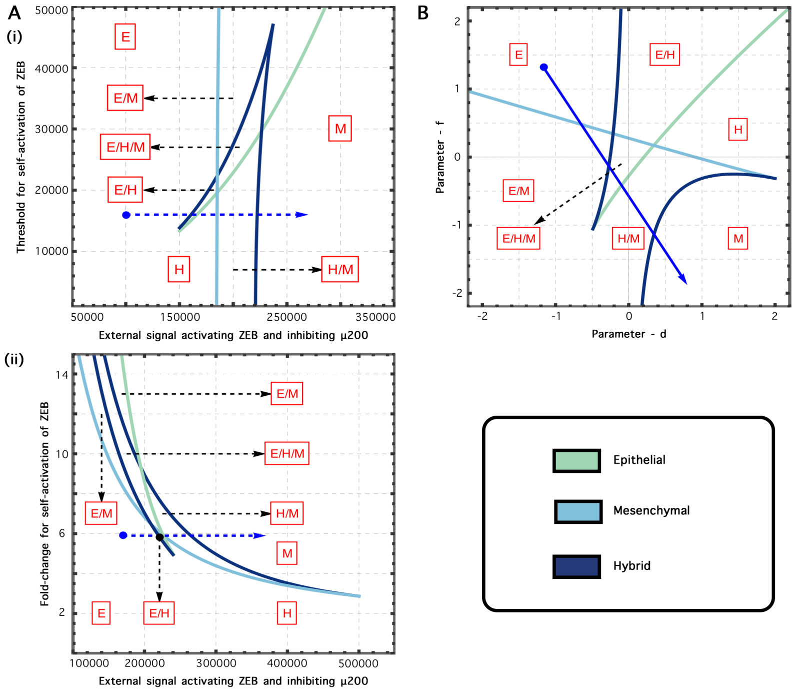


Figure 1: **Relating behaviour seen in ODE model to geometric model.** (A)(i) Bifurcation diagram of the threshold parameter of self-activation of ZEB, vs the external signal activating ZEB and inhibiting $\mu 200$. (ii) Bifurcation diagram of the fold-change parameter of self-activation of ZEB, vs the external signal activating ZEB and inhibiting $\mu 200$. (B) Bifurcation diagram of the two parameters - $d$ and $f$ in the butterfly equation which is analogous of behaviour of TSSA. Blue dashed lines in the three figures show a path following the same sequence of states (E→E/M→E/H/M→H/M→M).

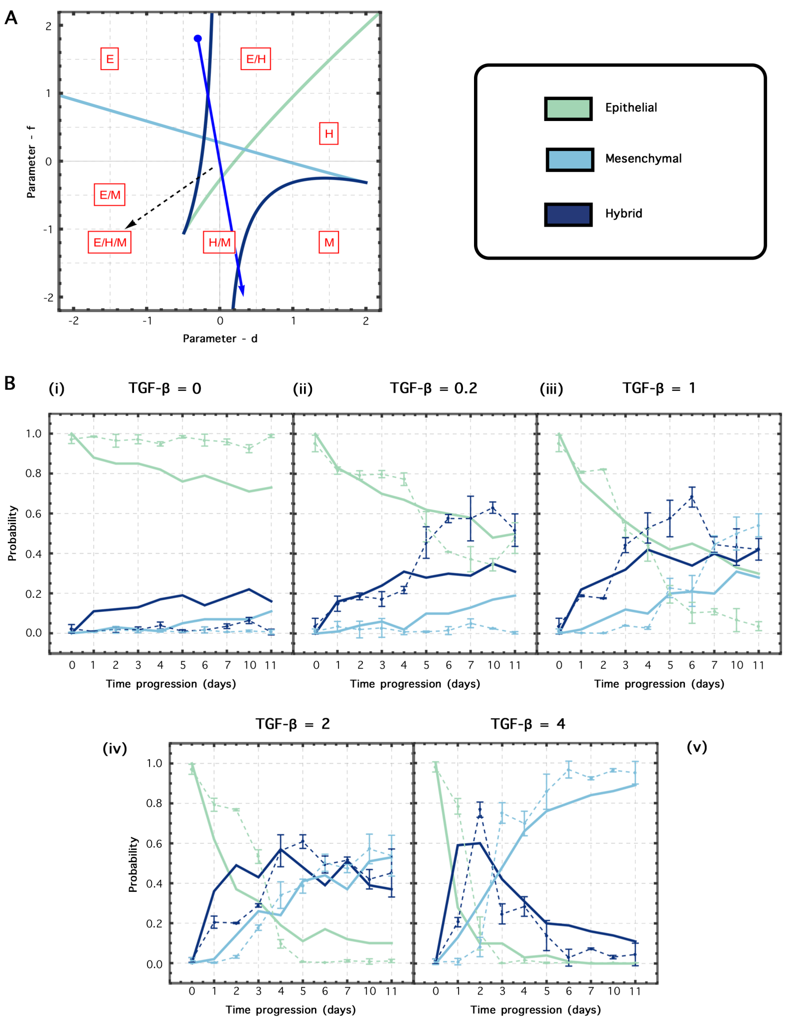


Figure 2: **Data fitting with another set of parameters.** (A) Bifurcation diagram of the two parameters - $d$ and $f$ in the butterfly equation which is analogous of behaviour of TSSA. (B)(i)-(v) Data corresponding to progression of population percentages of Epithelial, Mesenchymal and Hybrid populations at ten time points for different concentrations of TGF-$\beta$ (shown in dashed line). The population percentages as determined the fitted parameters (shown in solid line).
